# Cellular debris, not mucilage polysaccharides, dominate extracts of *Plantago ovata* and *Plantago lanceolata* calli grown *in vitro*

**DOI:** 10.1101/2020.06.15.153395

**Authors:** James M. Cowley, Lina Herliana, Natalie S. Betts, Rachel A. Burton

## Abstract

Mucilage is a hydrophilic mixture of polysaccharides produced by seeds of many species, and used in research, industrial processes and as human health supplements. As such, demand often outweighs supply. In recent years, several researchers have reported that mucilage can be produced efficiently from *in vitro-*cultured calli as a direct ‘plant-less’ alternative to seed mucilage; however, this mucilage has not been rigorously characterised. Here we replicate previously published culturing and extraction procedures and couple them with compositional analysis to determine whether the mucilage produced from *Plantago ovata* and *P. lanceolata* calli are similar to seed mucilage. Our monosaccharide profiling and microscopy show that, while calli derived from either seedling hypocotyls or roots yield more material than seeds using the same extraction techniques, the majority of extract mass is cellular debris. Debris polysaccharide composition was significantly different to seed mucilage from both species. Plant tissue culture as an alternative source of high yields of useful mucilage polysaccharides, at least by these and similar methods, is likely to not be possible.

## Introduction

Upon wetting, seeds of many plant species produce a sticky gel coating called mucilage. Mucilage consists primarily of highly hydrophilic polysaccharides, whose composition differs greatly between species^1^. Seed mucilage has research significance as a tool to study polysaccharide biosynthesis^2^, industrial significance as food hydrocolloids^3^, and health applications as viscous dietary fibres^4^. Demand for mucilage-producing seeds like psyllium (*Plantago ovata*) often exceeds supply due to agronomic constraints and large requirements in industrial and pharmaceutical applications, and so alternate sources of mucilage polysaccharides, such as *in vitro* production, is desirable.

Since early tissue culture experiments, increases in exogenous auxins were found to stimulate polysaccharide production^5–8^. Notably, calli of *Arabidopsis thaliana* stimulated with an exogenous auxin (2,4-dichlorophenoxyacetic acid; 2,4-D) would produce polysaccharides with a cell wall-like composition^8^. Undifferentiating calli producing polysaccharides with no assigned function or localisation—mucilage—are therefore referred to as mucilaginous calli (Figure 1). More recently, significant interest has been placed on mucilaginous calli as *ex planta* sources of commercial mucilage. It has been reported that mucilage produced from mucilaginous calli can be easily extracted to yield quantities significantly higher than seed-derived mucilage. Recent work reported that calli derived from *Plantago lanceolata*^9^, *P. ovata*^10,11^, *P. major*^12^, *Lepidium sativum*^13,14^, *Linum usitatissimum*^15^ and several *Alyssum* species^16^ could produce significantly (up to 10×) more mucilage than their respective seeds. As seed-derived mucilage from all of these species are important folk remedies, and that from *L. usitatissimum* and *Plantago* species have established industrial applications, this sort of efficient alternative production system would be of significant commercial interest. However, no studies have rigorously compared the composition and/or functional properties of callus-versus seed-derived mucilage to assess the feasibility of *in vitro* mucilage production. Due to our research group’s interest in seed mucilage, we were curious about the efficiency and possible applications of such methods so the aim of this work was to compare the composition of mucilage produced by calli to seed mucilage from two *Plantago* species.

**Figure 1.**
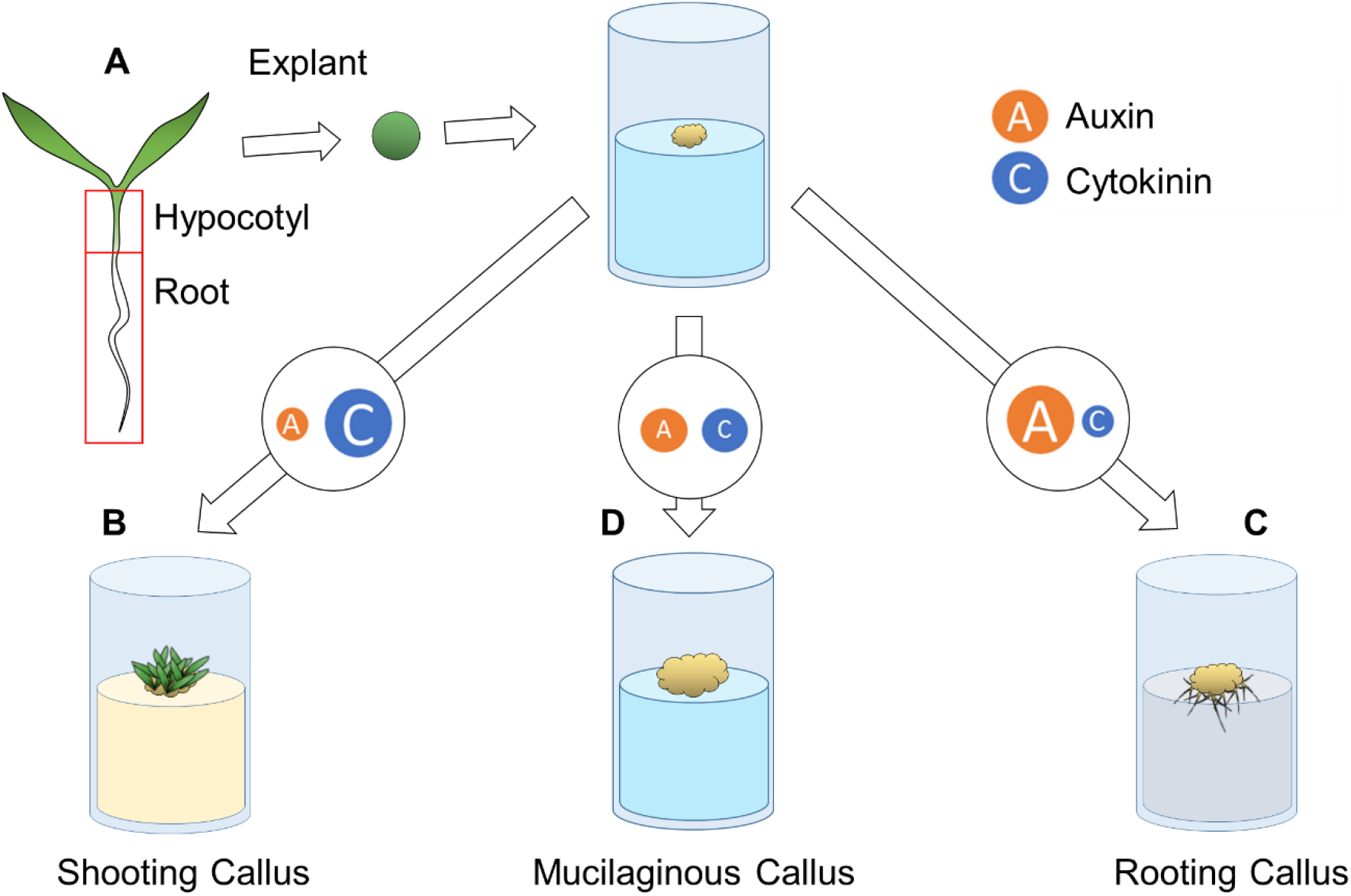
Varying levels of exogenous plant growth regulators can induce morphologically different calli *in vitro*. **A.** Pluripotent plant tissue explants (often hypocotyl or root) are used to induce callus. **B.** Low auxin:cytokinin favours shoot regeneration. **C.** High auxin:cytokinin favours root regeneration. **D.** When auxin and cytokinin are balanced, cells do not differentiate into distinct tissues and are referred to as mucilaginous calli.

## Results and Discussion

As a preliminary study, we performed tissue culture experiments on two seed mucilage-producing species established in our lab, *P. ovata* and *P. lanceolata.* These species were selected as they have similar germination times and conditions, and both produce heteroxylan-rich seed mucilage but with easily-distinguishable differences in polysaccharide fine structure^17^.

For optimisation of callogenesis, we opted for a 3×3 factorial arrangement of treatments (Table 1) using 2,4-D and kinetin (KIN) as the auxin and cytokinin, respectively, and using 10 day old hypocotyl and root tissues as explants. While the explant tissues of both species wilted rapidly in the absence of any hormones (T0 treatment), addition of any combination of hormones induced callogenesis (Figure 2A and 2B). Morphology of calli ranged from dark and compact to cream-coloured, soft and friable, and calli from *P. ovata* tended to be softer and wetter than that of *P. lanceolata*. Treatments T2 and T3 (lower 2,4-D to KIN) produced larger, softer, wetter-looking calli than other treatments; the dry weights of T2 and T3 calli were consistently highest across all treatments (Figure 2C), with T2 yielding the highest sum and grand mean for all replicates. The same treatment (0.5 mg/L 2,4-D with 1.0 mg/L KIN) had also been reported to produce very high callus induction rate, growth rate and mucilage yield from *P. ovata* hypocotyls^11^ and leaves^10^. Previous studies of mucilage production from *Plantago-*derived calli reported that the largest calli yielded the most mucilage^9,11^, so T2 calli from both species and explant types were used for mucilage compositional analysis.

**Table 1.**
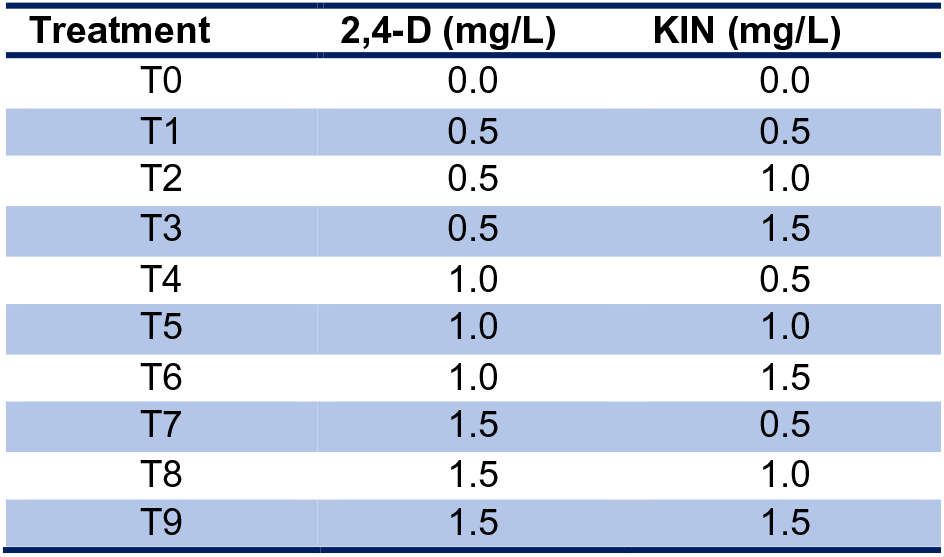
Hormone treatments.

| Treatment | 2,4-D (mg/L) | KIN (mg/L) |
| --- | --- | --- |
| T0 | 0.0 | 0.0 |
| T1 | 0.5 | 0.5 |
| T2 | 0.5 | 1.0 |
| T3 | 0.5 | 1.5 |
| T4 | 1.0 | 0.5 |
| T5 | 1.0 | 1.0 |
| T6 | 1.0 | 1.5 |
| T7 | 1.5 | 0.5 |
| T8 | 1.5 | 1.0 |
| T9 | 1.5 | 1.5 |

**Figure 2.**
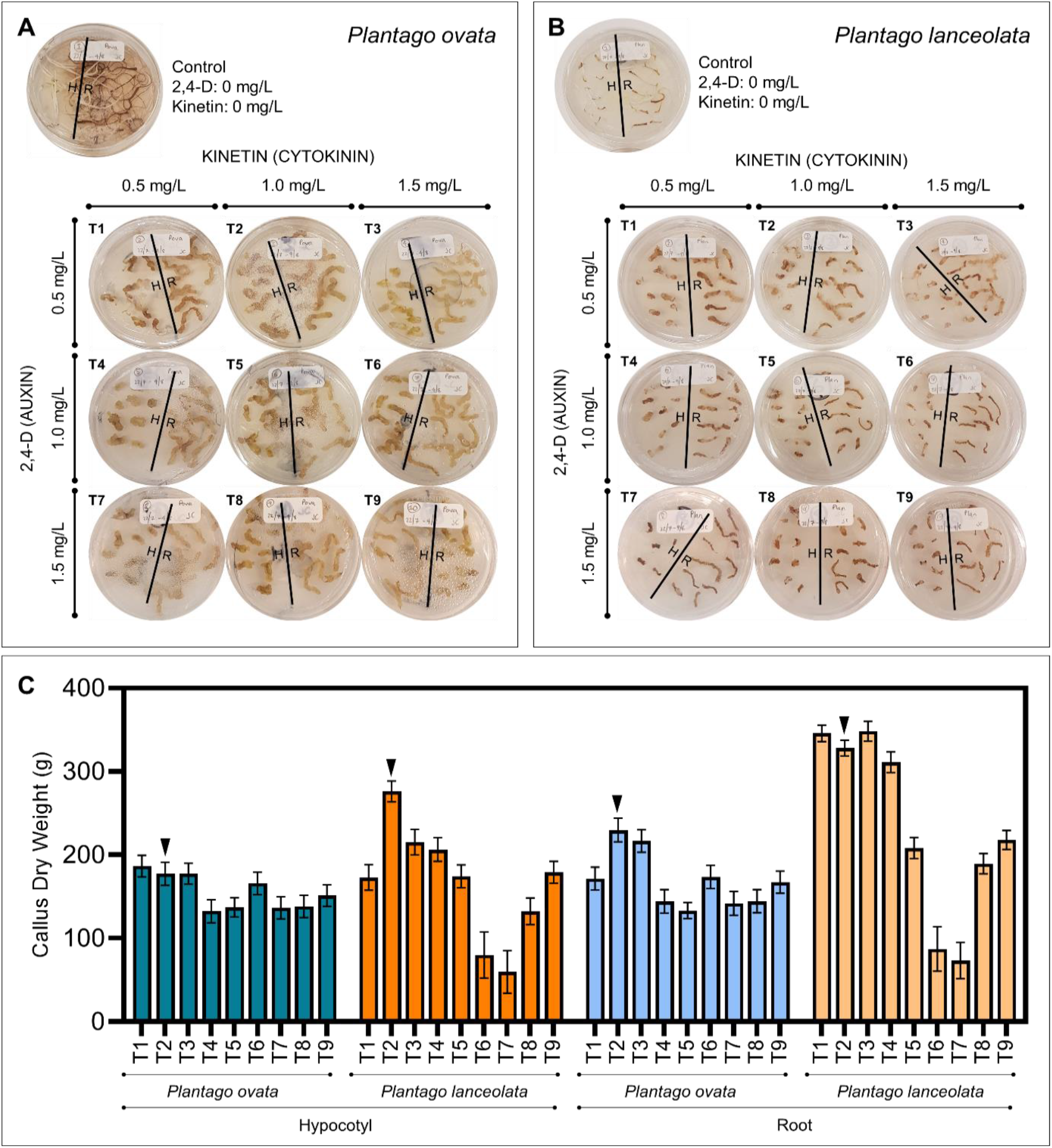
Effect of varying auxin and cytokinin ratios on *Plantago* callus morphology and yield. **A, B** Callus induction and morphology of (**A**) *P. ovata* and (**B)** *P. lanceolata* calli at 4 weeks. H, hypocotyl explant; R, root explant. **C.** Dry weight of calli after 8 weeks reveals that the effect of auxin and cytokinin are similar between species and explant type. Arrowhead denotes treatment 2 (T2), the highest yielding treatment that was used for composition analysis.

Mucilage was extracted from freeze-dried T2 calli alongside mature seeds. In line with previous findings^9,10,16^, the mass yield of material extracted from calli exceeded that extracted from seed in both species, but yield from both types of callus was comparable (Figure 3A). As mucilage is predominantly polysaccharide^1^, monosaccharide profiling was used to assess the composition of callus-derived extracted material (Figure 3B and 3C). While seed-derived mucilage from both *P. ovata* and *P. lanceolata* is clearly predominantly mono-or polysaccharide, material extracted from calli of both species had far more mass than could be accounted for as hydrolysed monosaccharides. While seed-derived mucilage is typically 70–90% polysaccharides^18,19^, only 15–30% of material extracted from calli could be recovered as monosaccharides after acid hydrolysis (Figure 3B). Comparing the molar monosaccharide composition of each sample, neither hypocotyl nor root callus-derived extracted material had a composition comparable to seed mucilage (Figure 3C). Monosaccharides present in callus-derived extracted material from both species were overwhelmingly glucose, compared with xylose-rich seed-derived mucilage (Figure 3C). As cellulose is resistant to the hydrolysis used here, the excess glucose is likely to be free cellular glucose or derived from starch. Though occurring in slightly different ratios, the remaining monosaccharides in the callus-derived extract corresponded to those in the post-extraction callus, which in turn corresponded to the composition of the fresh explant material (Figure 3C). The presence of arabinose, galactose and glucose were also reported for *L. sativum* callus mucilage^13^ but, similar to our results, these monosaccharides could only account for at most 20% of the mucilage mass. Furthermore, the molar ratios of these monosaccharides do not correspond to previous reports of the composition of *L. sativum* seed mucilage^20^. A similar compositional disconnect between hairy root culture-derived mucilage and gum of *Astragalus gummifer* has also been reported^21^. These findings indicate that callus-extracted monosaccharides are derived from non-cellulosic polysaccharides in the source cell walls, in line with some previous work^22^, rather than an extracellular mucilage.

**Figure 3.**
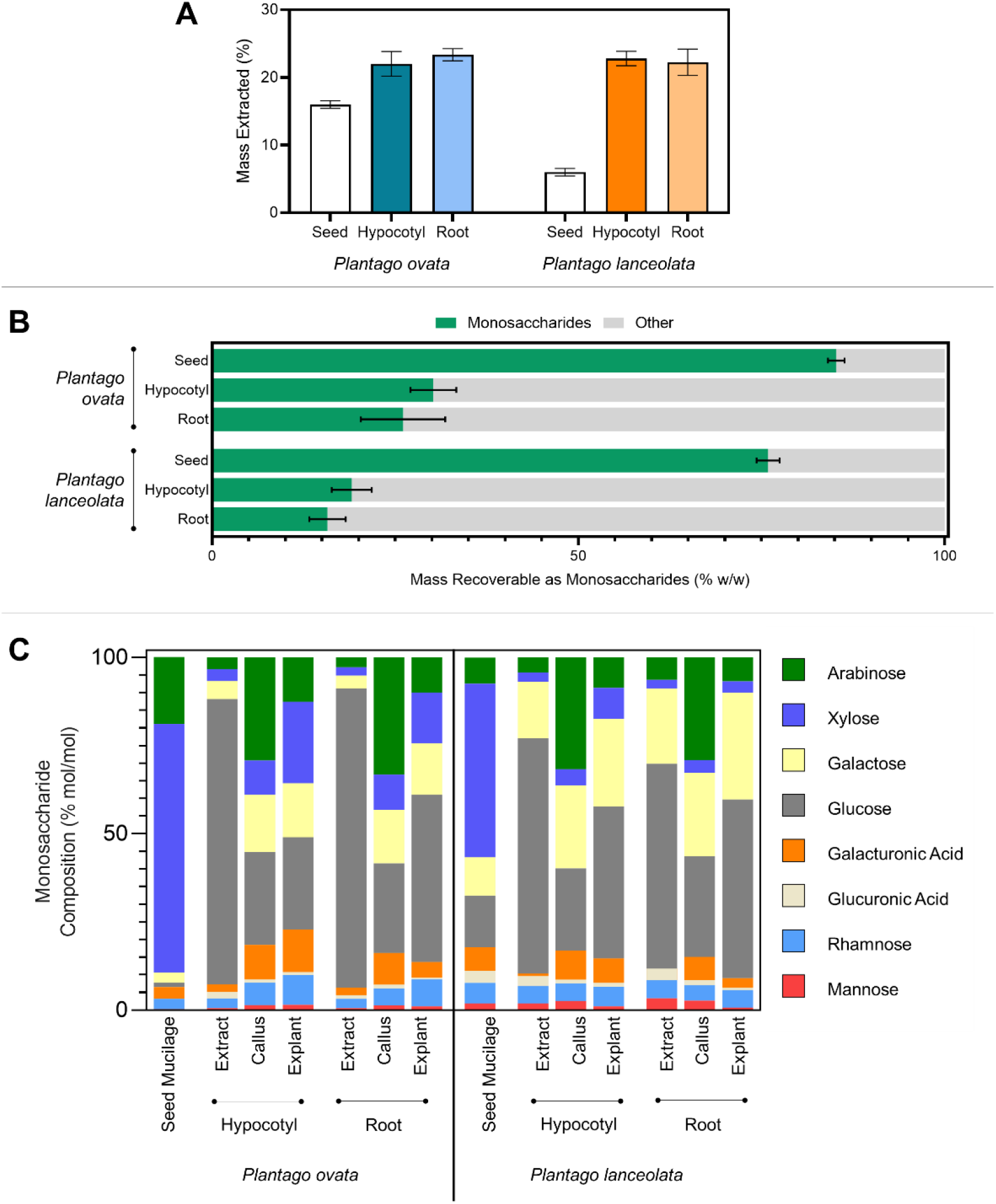
Yield and compositional analysis of *Plantago* callus extracts. **A.** Yield of extract from callus exceeds that of seed. **B.** Quantification of the proportion of extracts that could be recovered as monosaccharides. **C.** Molar monosaccharide profiles of extracts derived from various sources.

The callus extract mass that was not recoverable by monosaccharide analysis was substantial, and characterising the remaining material may shed light on why extract yields are, in agreement with findings here, reported to be so high. Based on the crude extraction techniques we hypothesised that the remaining mass may simply be insoluble debris. Using microscopic analysis we show that extracts from calli contain abundant particulate matter which is not present in corresponding extracts from seeds (Figure 4). Using the cellulose specific stain Direct Red 23^23^, it is clear that the debris is predominantly cellulosic wall material produced by the harsh extraction techniques and not effectively removed by filtering prior to further analysis. Even semi-intact, but somewhat damaged entire cells can be seen in *P. ovata* root callus extracts (Figure 4). Cellulosic materials are recalcitrant to hydrolysis by the technique used here, explaining the loss of mass recovery.

**Figure 4.**
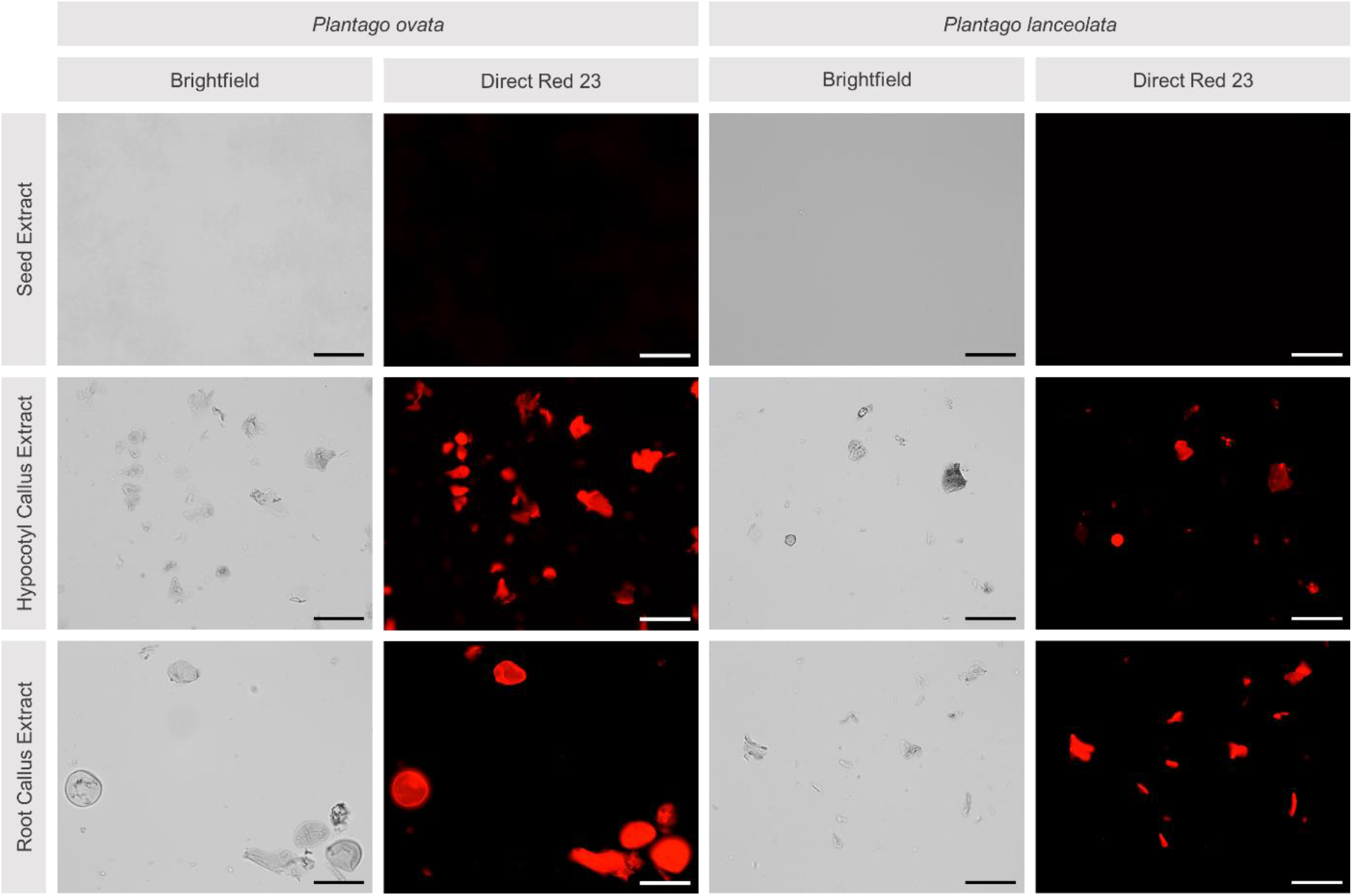
Microscopic analysis of *Plantago* seed and callus extracts. Cellular debris visible under brightfield stain positively with Direct Red 23. Scale bar = 100 µm.

## Conclusion

While previous reports of highly efficient *in vitro* mucilage production was an exciting premise, here we have shown that *Plantago* callus-derived extracted material, referred to by others as “mucilage”, is predominantly cellular debris, with a very different compositional profile to seed-derived mucilage from the same species. Previous reports of high yields are likely to be the result of ineffective filtering of cellular debris produced from harsh extraction techniques. Our monosaccharide analysis shows callus “mucilage” to have a low non-cellulosic polysaccharide content (only 21–35% of the content of seed mucilage), with an overall monosaccharide profile analogous to cell wall material from both callus and the source explant. We conclude that plant tissue culture as an alternative source of mucilage polysaccharides, at least by the methods followed here, is not likely to be possible.

## Methods

### Explant preparation and callus induction

*P. ovata* and *P. lanceolata* seeds were obtained from a previous study by Phan *et al.*^17^. Seeds were placed in an autoclaved stainless steel tea strainer and surface sterilised by sequential immersion for 1 min in filter-sterilised: 50% ethanol (once), 1:1 ethanol (50%) and bleach (4% hypochlorite) with 0.05% Triton X-100) (five times), and water (five times). Sterilised seeds were spread on sterile Whatman No. 1 filter paper in a Petri dish, wetted with 3 ml sterile water. Petri dishes were sealed with Parafilm, and seeds vernalised for 48 h at 4 °C before germination on a south-facing windowsill (Adelaide, Australia) for 10 days. Ten hypocotyl and ten root segments were aseptically excised from 10 day-old seedlings using a sterile scalpel and placed onto culture media (30 mg/L sucrose, 4.43 mg/L MS basal medium (PhytoTech, M519), 9% agarose) with different concentrations of 2,4-dichlorophenoxyacetic acid (2,4-D) (Astral Scientific, KB0745) and kinetin (KIN) (Astral Scientific, DB0166) (Table 1). Calli were incubated in the dark at 25 °C for 8 weeks, with subculturing every 2 weeks. Four replicates were prepared for each treatment and divided between two independent incubators.

At 8 weeks old, calli were harvested into 10 mL tubes, frozen at −80 °C and freeze-dried to a constant weight (approximately 72 hr)..

### Mucilage extraction

Freeze-dried 8 week old calli were lightly ground using a mortar and pestle prior to mucilage extraction. In line with previous publications^10,11,16^, mucilage was extracted from calli and seeds following the method of Sharma and Koul^24^ with some modifications described by Cowley *et al.*, ^25^. Briefly, 1.5 mL of milli-Q water was added to 50 mg of ground callus or seeds. Mixtures were extracted at 95°C for 30 minutes on a shaking incubator (Thermomixer Comfort, Eppendorf, Germany). Extracts were filtered through layered fine tulle (Spotlight, Australia) and freeze-dried for yield and monosaccharide analysis. This extraction was also tested with 0.1 M HCl (in place of water) as per Sharma and Koul^24^. No difference in yield or composition was observed, in line with our previous study^25^. As such, the hot water method was chosen for ease of handling and safety reasons.

### Monosaccharide analysis

Composition of mucilage extracted from seeds and callus, post-extraction callus material, and corresponding fresh explant material was determined by monosaccharide profiling following Cowley *et al.*,^25^.

### Microscopy

Freeze-dried extracts were dispersed at 2 mg/mL in 0.22 µm filtered milli-Q water. An aliquot of 967 µL was added to 33 µL of 5% w/v Direct Red 23 (Sigma-Aldrich) (1 in 30 dilution; final stain concentration of 0.165% w/v) and incubated in the dark at room temperature for 20 mins. One hundred microlitres of stain/extract was placed on a microscope slide and a cover slip placed on top.

Samples were imaged with a Zeiss M2 AxioImager with an AxioCam 506 mono black and white camera. Images were processed using ZEN 2012 software (Zeiss, Germany)

### Data analysis

Data analysis was performed in GraphPad Prism 8.4.0.

## Acknowledgements

This work was funded by the ARC Centre of Excellence in Plant Cell Walls (Grant No. 110001007). JMC is supported by a PhD scholarship from the Australian Government’s Research Training Program. The authors thank Rohan Singh for technical advice in plant tissue culture.

